# Reduced serotonin receptors and transporters in normal aging adults: a meta-analysis of PET and SPECT imaging studies

**DOI:** 10.1101/429266

**Authors:** Teresa M. Karrer, Casey L. McLaughlin, Carmela P. Guaglianone, Gregory R. Samanez-Larkin

**Author notes:** corresponding author: Teresa M. Karrer, M.Sc., Universitätsklinikum Aachen, Pauwelsstr. 30, 52074 Aachen, GERMANY.

## Abstract

Alterations in serotonin (5-HT) function have been hypothesized to underlie a range of physiological, emotional, and cognitive changes in older age. Here, we conducted a quantitative synthesis and comparison of the effects of age on 5-HT receptors and transporters from cross-sectional PET and SPECT imaging studies. Random-effects meta-analyses of 31 studies including 1087 healthy adults yielded large negative effects of age in 5-HT-2A receptors (largest in global cortex), moderate negative effects of age in 5-HT transporters (largest in thalamus), and small negative effects of age in 5-HT-1A receptors (largest in parietal cortex). Presynaptic 5-HT-1A autoreceptors in raphe/midbrain, however, were preserved across adulthood. Adult age differences were significantly larger in 5-HT-2A receptors compared to 5-HT-1A receptors. A meta-regression showed that 5-HT target, radionuclide, and publication year significantly moderated the age effects. The findings overall identify reduced serotonergic signal transmission in healthy aging. The evidence for the relative preservation of 5-HT-1A compared to 5-HT-2A receptors may partially explain psychological age differences, such as why older adults use more emotion-focused rather than problem-focused coping strategies.

## 1. Introduction

Serotonin (5-HT) is involved in a wide range of physiological, emotional, and cognitive functions (Lucki, 1998). The existence of at least 14 distinct receptor types that differ in their distribution across the brain might explain the functional diversity of the 5-HT system (Barnes and Sharp, 1999). Although the majority of studies of the functional consequences of alterations in the 5-HT system have focused on the role of 5-HT transporters in depression (Gryglewski et al., 2014; Kambeitz and Howes, 2015; Ogilvie et al., 1996), 5-HT receptors have been specifically associated with several functions. For example, the inhibitory 5-HT-1A receptor has been linked to the sleep-wake cycle, the modulation of anxiety, and long-term memory, whereas the excitatory 5-HT-2A receptor has been associated with the modulation of hallucinogenic effects and both short and long-term memory (Gaspar et al., 2003; Hannon and Hoyer, 2008). Moreover, 5-HT dysfunctions in general have been implicated in several psychiatric and neurological diseases across the life span (Lucki, 1998) including age-related diseases like Alzheimer’s disease (Blin et al 1993).

In contrast to candidate gene studies which make up the majority of individual difference studies of 5-HT in humans (Caspi et al., 2010), positron emission tomography (PET) and single-photon emission computed tomography (SPECT) are brain imaging techniques that provide direct measures of 5-HT system function in-vivo. Over the last several decades, one of the most consistently observed findings in molecular imaging studies of human brain 5-HT targets is a decline with healthy aging. Studies of different receptor types and transporters have revealed small to moderate negative effects of age on 5-HT-1A receptors (Rabiner et al., 2002; Tauscher et al., 2001), moderate to large effects of age on 5-HT-2A receptors (Sheline et al., 2002; Uchida et al., 2011), and moderate to large effects of age on 5-HT transporters (Fazio et al., 2016; Yamamoto et al., 2002) across adulthood. The only prior summary of the literature on 5-HT in aging, based primarily on post-mortem studies, concluded that 5-HT-1A and 5-HT-2A receptors decline with age (Meltzer et al., 1998). The evidence for adult age differences in 5-HT transporters was mixed, including findings of reduced, increased, and stable 5-HT transporter levels with age (Meltzer et al., 1998).

From neuroimaging studies, the previously reported sizes of adult age effects varied across 5-HT targets and cortical and subcortical brain regions. Some of the observed heterogeneity might be due also to sample variation in other variables of influence such as sex (Cidis Meltzer et al., 2001; Costes et al., 2005), body mass index (Hesse et al., 2014), hormone levels (Moses-Kolko et al., 2011), and genes (David et al., 2005). It is also possible that different 5-HT targets do not change uniformly across the life span but instead develop and age in a non-linear way. This might produce different estimates of age effects depending on the specific age ranges within each study. Furthermore, many PET and SPECT imaging studies lack appropriate sample sizes due to the high costs and safety concerns of nuclear imaging technology. Thus, it is difficult to assess the representativeness or accuracy of a single effect from most PET and SPECT studies, as they usually include less than 25 participants per group.

Recent meta-analytic approaches have been used to aggregate and compare individual effects across studies. Although decades of research have documented age-related declines in neuromodulatory function in general, recent meta-analytic approaches have identified differential rates of decline across subcomponents of neuromodulatory systems. For example, a quantitative synthesis of adult age effects on the dopamine system in over 2000 human participants revealed moderate to large negative effects of age on D1- and D2-like receptors and dopamine transporters (Karrer et al., 2017). Critically, the age effect in D1-like receptors was significantly larger than in D2-like receptors, whereas presynaptic synthesis capacity was relatively preserved across adulthood (Karrer et al., 2017). Although a qualitative summary of the existing neuroimaging literature suggests moderate to strong negative effects of age across the 5-HT system, human neuroimaging studies have not systematically and directly compared effects of age across different 5-HT targets. Here, we used a meta-analytic approach to investigate variation in the effects of aging across the 5-HT system.

To quantitatively synthesize the existing literature on adult age differences in several targets of the 5-HT system, we conducted a comprehensive meta-analysis of cross-sectional PET and SPECT imaging studies. In light of previous individual study findings, we expected small to moderate negative effects of age on 5-HT-1A receptors and moderate to large negative effects of age on 5-HT-2A receptors and transporters. To investigate which factors might explain the heterogeneity of findings across previous studies, we examined the influence of potential moderating variables (e.g., age range, sex, imaging method, 5-HT target, brain region) in a meta-regression. In additional more exploratory analyses, we aggregated individual participant data across studies (where available) to examine the linearity of age effects across 5-HT targets. We did not make specific predictions for these exploratory analyses given the lack of prior reporting and low power of individual studies to examine these effects.

## 2. Methods

We conducted and reported this meta-analysis adhering to the guidelines of the PRISMA (Preferred Reporting Items for Systematic Reviews and Meta-Analyses) statement (Liberarti et al., 2009). Note that we did not pre-specify our approach in a review protocol. Instead, we closely followed the methods described in an earlier meta-analysis of our group on age differences in the dopamine system (Karrer et al., 2017). In addition, this facilitates the comparison of age effects across both neurochemical systems.

### 2.1 Literature search

Potentially relevant studies from the personal library of G.R.S.-L. helped to define the keywords for the systematic literature search. We capitalized on the hierarchically organized vocabulary of Medical Subject Headings (MeSH) terms that enables a search for specific topics independent of the used words or spellings (Lipscomp, 2000). The MeSH terms “aging”, “emission computed tomography”, “humans”, “serotonin receptors”, “serotonin plasma membrane transport proteins”, “serotonin” and “synthesis” were combined to search the largest biomedical literature database PubMed (cf. Supplementary Materials for exact literature search terms). The last search was performed on 01/16/2019. To further complement the study corpus, we carefully checked the references of all relevant studies to search for additional includable studies.

We included all accessible PET and SPECT imaging studies of the 5-HT system written in English or German that (i) reported original results, (ii) contained a sample of healthy adults (> 18 years) with a minimum age range of 25 years to representatively measure the age effect, and (iii) reported or allowed for the computation of the Pearson correlation coefficient *r* between age and a 5-HT target. We included different tracer kinetic measures that quantify the functioning of 5-HT targets such as non-displaceable binding potential and binding potential based on an arterial input function (Innis et al., 2007). We excluded studies (i) if their sample primarily consisted of relatives of depressive patients because serotonergic alterations have been observed in this subgroup (Leboyer et al., 1999), (ii) if we could not extract a standardized main effect of age comparable to Pearson’s *r* (e.g., in multiple regressions), and (iii) if their sample could be identified as subsample of another included study. To avoid duplicates in the included participants, we thoroughly inspected sample descriptions (e.g., age range and female percent) and compared experimental characteristics (e.g., scanner model and radiotracer) across studies.

### 2.2 Data extraction

Given that many studies reported the Pearson correlation coefficient *r* between age and individual targets of the 5-HT system (receptors, transporters, synthesis capacity), we used this standardized statistic in all meta-analytic analyses. The included studies either directly reported or enabled the computation of the Pearson correlation coefficient *r* from (i) a table, (ii) a figure (using PlotDigitizer version 2.6.8), or (iii) the group effect of age in older versus younger adults. We summarized the age effects for each of the following brain regions: raphe/midbrain, thalamus, medial temporal lobe (amygdala, hippocampus, parahippocampal gyrus, entorhinal cortex, and perirhinal cortex), cingulate, insula, occipital, parietal, temporal, frontal and global cortex. Note that in case of 5-HT-1A receptors, raphe/midbrain regions predominantly contain presynaptic autoreceptors as opposed to the postsynaptic receptors that are in terminal regions (Barnes and Sharp, 1999). If a study reported the age effect separately for males and females, right and left hemisphere, or several sub-regions of the brain, we averaged the correlation coefficients using Fisher’s *z* transformation. Particularly in small samples, this conversion to an approximately normally distributed measure introduces less bias in the computation of the mean (Silver and Dunlap, 1987). If a study computed the age effects using several nuclear imaging analysis methods, we chose the method which the authors referred to as gold-standard or which was most commonly used in the literature.

In addition to the effect sizes of primary interest (*r* between 5-HT target and age), we extracted the following data: sample size, age range, female percent, imaging method, scanner model, 5-HT target, brain region, radiotracer, tracer kinetic measure, reference region, and partial volume correction. To examine the nature of the age effects in more detail, we extracted age and 5-HT target data of individual participants from tables or figures reported in a majority of the included studies. All extracted data are available at osf.io/3897j.

### 2.3 Data analysis

First, we examined the possibility that studies reporting stronger effects were more likely to get published and thus, included in the meta-analyses. To qualitatively assess publication bias, we used contour-enhanced funnel plots. In contrast to the traditional version, this visual tool indicates areas of statistical significance. This approach distinguishes publication bias from other types of biases such as study quality (Peters et al., 2008). We additionally applied the rank correlation test (Begg and Mazumdar, 1994) to quantitatively test for publication bias.

The primary goal was to meta-analytically summarize the correlations between age and different 5-HT targets across studies. We estimated one meta-analysis for each 5-HT target split into several target regions. Because the studies were based on different populations and varied in many aspects, we chose random-effects models to account for the heterogeneity between studies (Borenstein et al., 2010). In case the meta-analyses relied on too few studies to robustly estimate the heterogeneity between studies, we additionally compared the results to fixed-effect models, which only assume variance within the studies (Borenstein et al., 2010). If a study provided age correlations for more than one region, we used a multi-variate meta-analysis model to account for the statistical dependence of the effect sizes (Jackson et al., 2011). To stabilize the variances of the effect sizes, Pearson’s correlation coefficients *r* were converted to Fisher’s *z* before analysis (e.g., Viechtbauer, 2010).

In the next step, we examined how much of the overall study heterogeneity could be attributed to differences between studies rather than errors within studies using *Q* and *I^2^* statistics (Higgins et al., 2003). To investigate which variables might explain the observed heterogeneity between the studies, we used a mixed-effects meta-regression. We included the following potential moderator variables in the meta-regression: age range and female percent of the sample, publication year, imaging method, radionuclide, tracer kinetic measure, reference region, partial volume correction, 5-HT target, and brain region (Table S1). We centered continuous variables first to facilitate interpretation. Since some studies contributed several effect sizes to the meta-regression, we again used a multivariate model that accounts for the statistical dependence of the effect sizes.

Because the study-level meta-analyses of linear effects of age were limited in providing insight into potential nonlinear associations between age and 5-HT targets, we additionally analyzed individual participant data. We estimated the type of relationship between age and different 5-HT targets using raw data extracted from a subset of the studies. If a study reported the tracer kinetic measure values for several brain regions instead of a global aggregation, we averaged the tracer kinetic measure values in each participant first. Then, we z-scored the individual tracer kinetic measure values in each study to allow for a comparison across different tracer kinetic measures. Finally, we fit linear, logarithmic, and quadratic functions to the individual data points to investigate which model best described the relationship between age and 5-HT targets. Mechanistically, linear models imply zero-order processes (e.g., constant decline per time unit), whereas logarithmic models suggest first-order processes (e.g., amount of decline dependent on 5-HT target concentration). Quadratic models, however, might indicate several competing processes. The extracted raw data additionally enabled us to estimate the percent difference per decade for different 5-HT targets. For this purpose, we divided the slope of the linear function by the range of the tracer kinetic measure values and multiplied the proportion by 10.

Finally, the meta-analytically-derived summary effect sizes were used to conduct statistical power analyses. Based on a two-sided significance level of *alpha* = 0.05, we determined the percentage of included studies with at least 80% statistical power. In addition, we computed the necessary sample sizes to detect effects of age in different 5-HT targets, which could be useful for future studies.

Analyses were conducted in R (version 3.4.0) primarily using the metafor package (Viechtbauer, 2010). All data analysis scripts are publicly available at osf.io/ctn83 for transparency and reuse.

## 3. Results

### 3.1 Literature search

The literature search yielded 299 potentially relevant studies. The study selection process is detailed in a PRISMA flowchart (Figure S1). Studies reporting age effects in 5-HT-1B receptors (Matuskey et al., 2012; Nord et al., 2014), 5-HT-4 receptors (Madsen et al., 2011; Marner et al., 2010), and 5-HT synthesis capacity (Rosa-Neto et al., 2007) were excluded due to an insufficient number to conduct a meta-analysis. A total of 31 studies reporting age effects in 5-HT-1A receptors (n=8), 5-HT-2A receptors (n=18), and 5-HT transporters (n=6) were included in the meta-analyses (cf. Supplementary Materials for the reference list; one study examined age effects in two 5-HT targets (Moses-Kolko et al., 2011)). The study corpus included 83.9% PET studies, 35.1% females, and a mean age range of 49 years in 1087 healthy adults (Table S2).

### 3.2 Publication bias and meta-analysis

The funnel plots contained few non-significant studies in 5-HT-2A receptors and 5-HT transporters, which could indicate publication bias (Figure S2). However, the applied rank correlation tests did not detect any significant relationship between effect sizes and sampling variance in 5-HT-2A receptors (*Kendall’s τ* = −0.18*; p* = 0.149) and in 5-HT transporters (*Kendall’s τ* = *−0.15; p* = 0.615). In 5-HT-1A receptors, however, the rank correlation test yielded a significant funnel plot asymmetry (*Kendall’s τ* = −0.30*; p* = 0.010). The random-effects meta-analyses estimated the summary correlation coefficients and their 95% confidence intervals. The average age effect was *r* = –0.23 [–0.37; –0.08] in 5-HT-1A receptors (Figure 1), *r* = –0.71 [–0.77; –0.63] in 5-HT-2A receptors (Figure 2), and *r* = −0.49 [−0.65; −0.28] in 5-HT transporters (Figure 3). Different variants of univariate and fixed-effect meta-analytic models yielded comparable results (Table S3). In addition, we computed the summary effect sizes separately for different brain regions (Table 1). We consistently observed negative effects of age across 5-HT target and brain region with the exception of 5-HT-1A receptors in raphe/midbrain and cingulate, which did not show significant age-related differences. The comparison of the 95% confidence intervals revealed significantly larger age differences in 5-HT-2A receptors compared to 5-HT-1A receptors (Figure 4).

**Figure 1.**
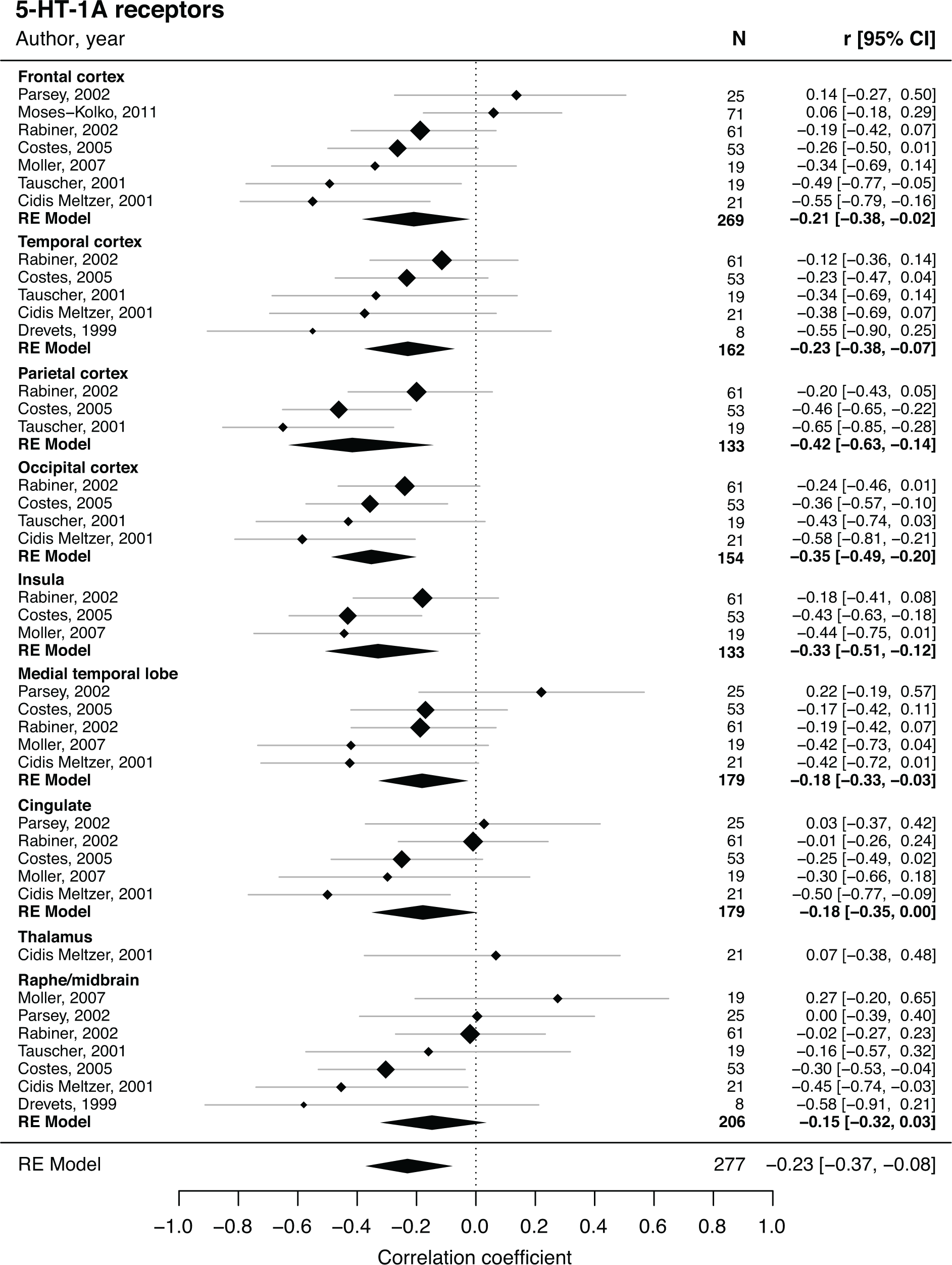
Forest plot of age effects in 5-HT-1A receptors. Depicts reported effect size (diamond) and 95% confidence interval (line) for each included study. Weight of each study (size of diamond) depends on sample size and between-study variance. Summary effects sizes (polygon) and 95% confidence interval (width of polygon) for separate brain regions and in total. N represents the number of individual participants included in each study and in the computation of the summary effect sizes.

**Figure 2.**
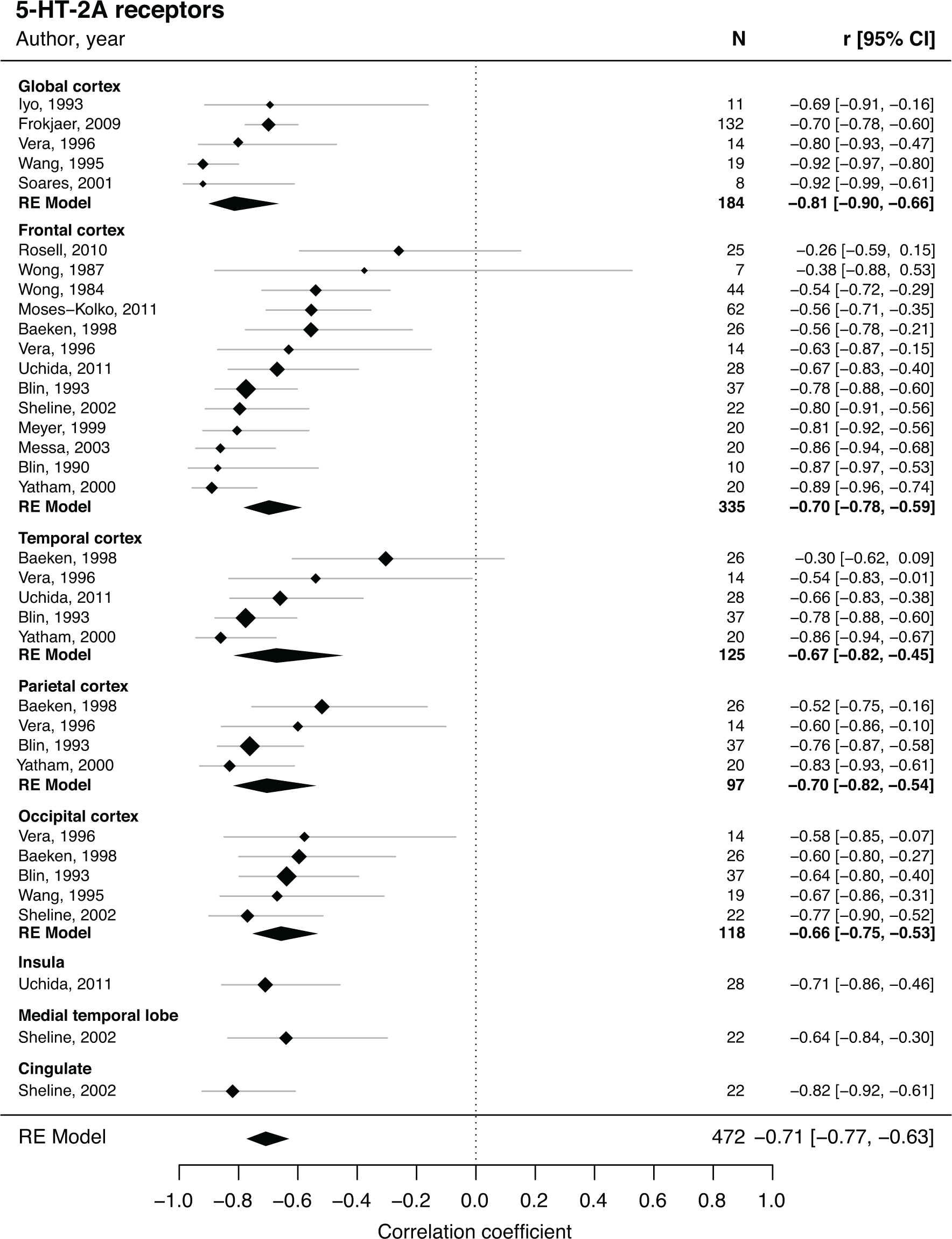
Forest plot of age effects in 5-HT-2A receptors. Depicts reported effect size (diamond) and 95% confidence interval (line) for each included study. Weight of each study (size of diamond) depends on sample size and between-study variance. Summary effects sizes (polygon) and 95% confidence interval (width of polygon) for separate brain regions and in total. N represents the number of individual participants included in each study and in the computation of the summary effect sizes.

**Figure 3.**
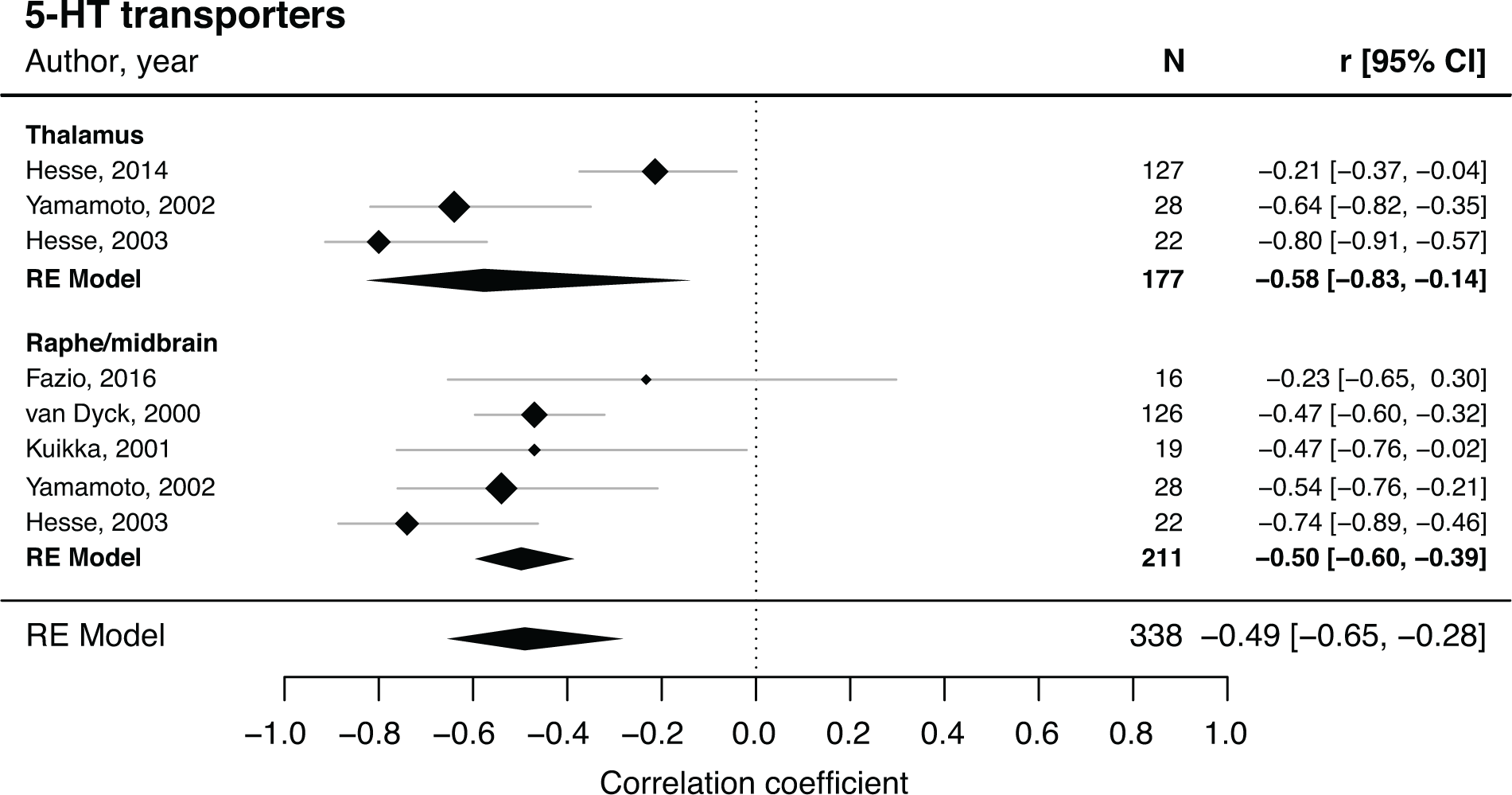
Forest plot of age effects in 5-HT transporters. Depicts reported effect size (diamond) and 95% confidence interval (line) for each included study. Weight of each study (size of diamond) depends on sample size and between-study variance. Summary effects sizes (polygon) and 95% confidence interval (width of polygon) for separate brain regions and in total. N represents the number of individual participants included in each study and in the computation of the summary effect sizes. Note that in one study (Hesse et al., 2003) the region of interest for thalamus also included hypothalamus.

**Figure 4.**
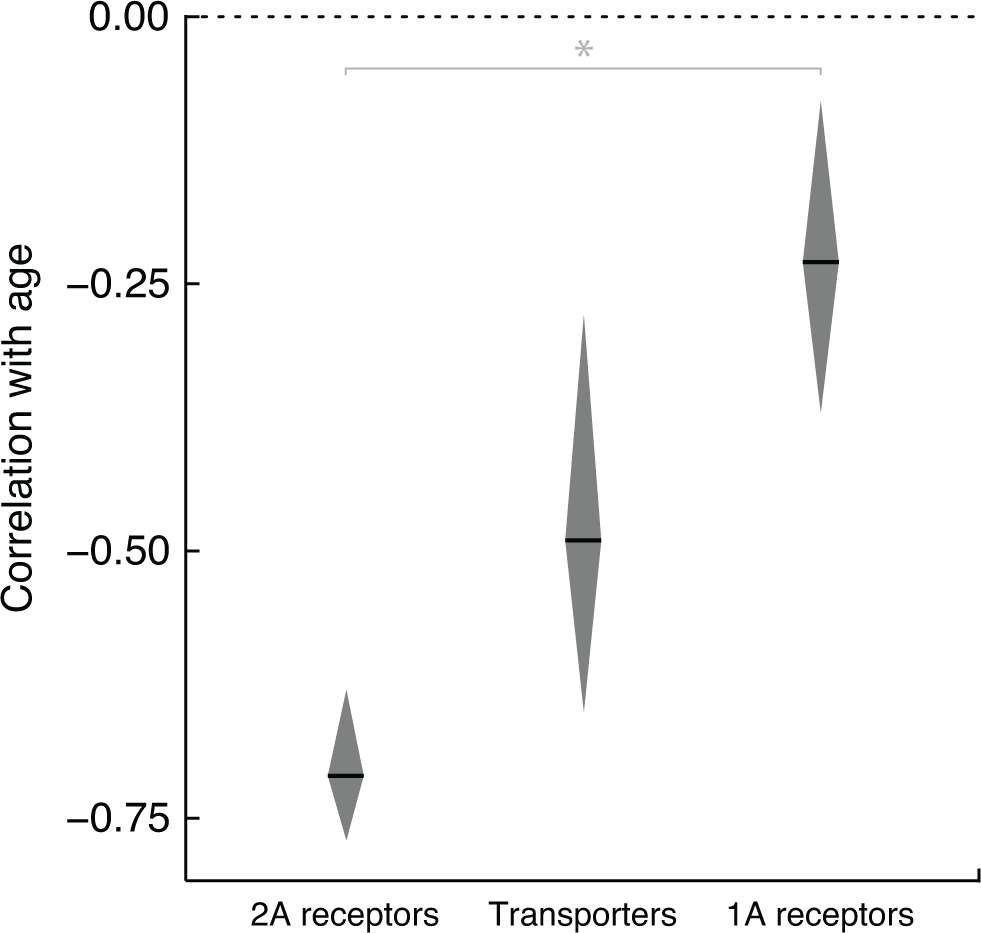
Comparison of overall age effects in 5-HT targets. Effects compared here are based on the lowermost meta-analytically-derived summary effect size polygons from Figures 1–3 in 5-HT-1A receptors (across frontal, temporal, parietal, and occipital cortex, insula, medial temporal lobe, cingulate, and raphe/midbrain), 5-HT-2A receptors (across global, frontal, temporal, parietal, and occipital cortex), and 5-HT transporters (across thalamus and raphe/midbrain). Black line indicates summary effect size and height of polygon represents 95% confidence interval. Dotted line marks an age effect of zero. Age effects on 5-HT-1A receptors and 5-HT-2A receptors were significantly different at *p < 0.01 (Cumming, 2009).

**Table 1.** Results of meta-analyses of adult age differences in 5-HT-1A receptors, 5-HT-2A receptors, and 5-HT transporters.

| Target | r | 95% CI |  | # of studies | # of participants |
| --- | --- | --- | --- | --- | --- |
| **5-HT-1A receptor** | -0.23 | -0.37 | -0.08 | 8 | 277 |
| Frontal cortex | -0.21 | -0.38 | -0.02 | 7 | 269 |
| Temporal cortex | -0.23 | -0.38 | -0.07 | 5 | 162 |
| Parietal cortex | -0.42 | -0.63 | -0.14 | 3 | 133 |
| Occipital cortex | -0.35 | -0.49 | -0.2 | 4 | 154 |
| Insula | -0.33 | -0.51 | -0.12 | 3 | 133 |
| Medial temporal lobe | -0.18 | -0.33 | -0.03 | 5 | 179 |
| Cingulate | -0.18 | -0.35 | 0 | 5 | 179 |
| Raphe/midbrain | -0.15 | -0.32 | 0.03 | 7 | 206 |
| **5-HT-2A receptor** | -0.71 | -0.77 | -0.63 | 17 | 472 |
| Global cortex | -0.81 | -0.9 | -0.66 | 5 | 184 |
| Frontal cortex | -0.7 | -0.78 | -0.59 | 13 | 335 |
| Temporal cortex | -0.67 | -0.82 | -0.45 | 5 | 125 |
| Parietal cortex | -0.7 | -0.82 | -0.54 | 4 | 97 |
| Occipital cortex | -0.66 | -0.75 | -0.53 | 5 | 118 |
| **5-HT transporter** | -0.49 | -0.65 | -0.28 | 6 | 338 |
| Thalamus | -0.58 | -0.83 | -0.14 | 3 | 177 |
| Raphe/midbrain | -0.5 | -0.6 | -0.39 | 5 | 211 |

### 3.3 Heterogeneity and meta-regression

Significant *Q*-statistics in 5-HT-1A receptors (*Q* = 58.0%; *p* = 0.026), 5-HT-2A receptors (*Q* = 61.6%; *p* = 0.003), and 5-HT transporters (*Q* = 23.2%; *p* = 0.002) indicated that the included studies did not share a common effect size. Similarly, the *I^2^*-statistics revealed moderate amounts of heterogeneity in 5-HT-1A receptors (*I^2^* = 55.1%), 5-HT-2A receptors (*I^2^* = 55.9%), and 5-HT transporters (*I^2^* = 72.5%). The mixed-effects meta-regression showed that the included variables explained a significant amount of the observed heterogeneity in the age effect (*p* < 0.001). Publication year (*p* = 0.005) and radionuclide (*p* = 0.008) emerged as significant moderators of the age effect (Table 2). Specifically, smaller effects of age were reported in more recent publications and in studies using the radiotracer ^11^C instead of ^18^F. Post-hoc tests, which evaluated the influence of a variable across several factor levels, additionally showed a significant moderating effects of 5-HT target (*p* = 0.001). Age range, female percent, imaging method, tracer kinetic measure, reference region, partial volume correction, and brain region did not significantly moderate the age effect. A test for residual heterogeneity indicated that other variables not considered in the meta-regression did not significantly influence the age effect (*p* = 0.226).

**Table 2.** Influence of potential moderators on the correlation between age and 5-HT targets in a multi-variate mixed-effects meta-regression.

| Moderator variable<br>(factor level) | r | SE | 95% CI |  |
| --- | --- | --- | --- | --- |
| Intercept | -0.13 | 0.12 | -0.35 | 0.11 |
| Age range | 0 | 0.01 | -0.01 | 0.01 |
| Female percent | -0.25 | 0.31 | -0.69 | 0.33 |
| Publication year | 0.02 | 0.01 | 0.01 | 0.04 |
| Imaging method (SPECT) | 0.18 | 0.23 | -0.27 | 0.56 |
| Tracer kinetic measure (BP<sub>ai</sub>) | 0.24 | 0.13 | -0.01 | 0.45 |
| Radionuclide (<sup>18</sup>F) | -0.39 | 0.16 | -0.62 | -0.11 |
| Reference region (occipital) | 0.24 | 0.25 | -0.24 | 0.62 |
| Partial volume correction (Applied) | -0.01 | 0.14 | -0.28 | 0.25 |
| Target (5-HT transporter) | -0.66 | 0.22 | -0.84 | -0.35 |
| Target (5-HT-2A receptor) | -0.3 | 0.15 | -0.55 | -0.01 |
| Region (Cingulate) | -0.12 | 0.1 | -0.31 | 0.09 |
| Region (Frontal cortex) | -0.19 | 0.09 | -0.36 | 0 |
| Region (Global cortex) | -0.54 | 0.21 | -0.77 | -0.2 |
| Region (Insula) | -0.24 | 0.11 | -0.43 | -0.02 |
| Region (Medial temporal lobe) | -0.09 | 0.1 | -0.29 | 0.11 |
| Region (Occipital cortex) | -0.23 | 0.1 | -0.41 | -0.02 |
| Region (Parietal cortex) | -0.25 | 0.11 | -0.44 | -0.04 |
| Region (Temporal cortex) | -0.09 | 0.1 | -0.28 | 0.11 |
| Region (Thalamus) | 0 | 0.14 | -0.27 | 0.27 |

*Note:* Three studies missing information were automatically dropped from the meta-regression. The factor level “<sup>123</sup>I” of the radionuclide variable was automatically dropped from the model due to a complete redundancy with the factor level “SPECT” of the imaging method variable.

### 3.4 Linearity/non-linearity of age effects and percent difference per decade

Individual participant data for at least a subset of brain regions were available for nearly all of the studies (n=29 out of 31). This allowed us to test linear, logarithmic, and quadratic effects of age across 5-HT targets (Figure 5). Linear age effects explained 2% of the variance in 5-HT-1A receptors (*p* < 0.05), 41.7% of variance in 5-HT-2A receptors (*p* < 0.001) and 11.7% of variance in 5-HT transporters (*p* < 0.001). The fits of the logarithmic and quadratic functions were comparable to those of the linear function while adjusting for model complexity (Table S4). The quadratic term, however, did not add a significant amount of explained variance in any of the three 5-HT targets (*p_5-HT-1A_*= 0.976; *p_5-HT-2A_* = 0.727; *p_5-HT_* = 0.934).

**Figure 5.**
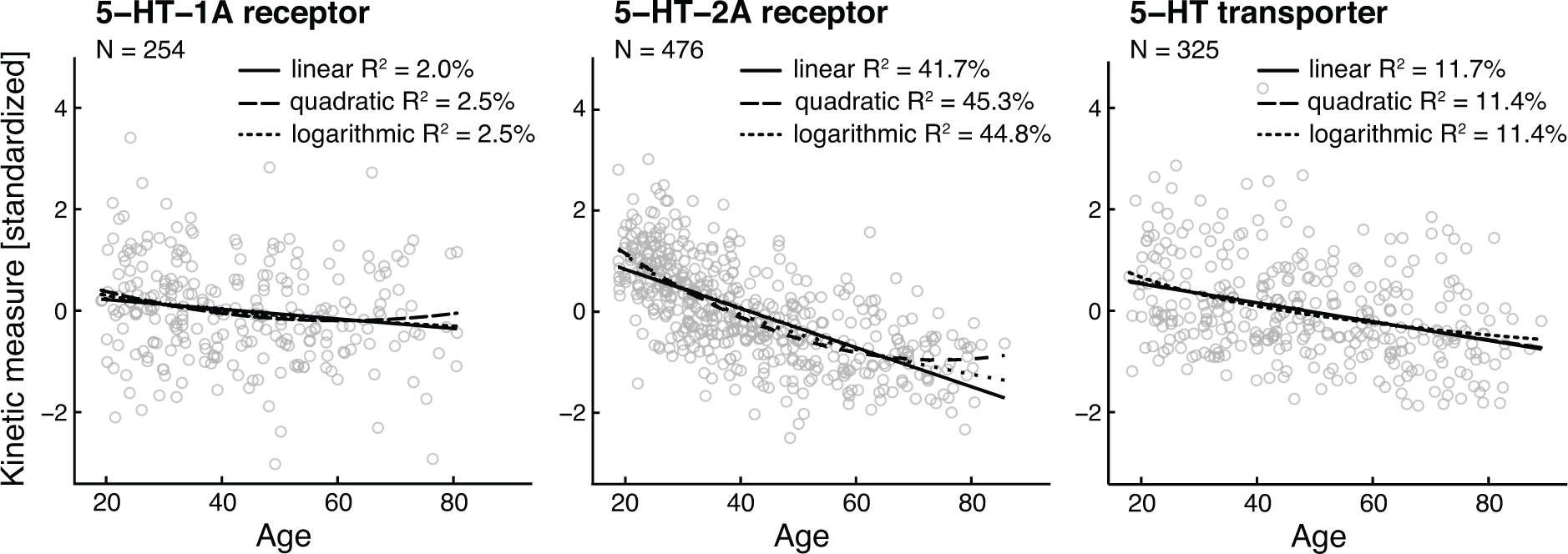
Scatter plots depicting the relationship between age and 5-HT targets based on a subset of studies (29 out of 31 studies) for which raw data were available. Data points represent individual age effects averaged across brain regions for 5-HT-1A receptors (across frontal, temporal, parietal, and occipital cortex, insula, medial temporal lobe, cingulate, and raphe/midbrain), 5-HT-2A receptors (across global, frontal, temporal, parietal, and occipital cortex), and 5-HT transporters (across thalamus and raphe/midbrain). Linear, quadratic, and logarithmic model were fit to individual age and standardized tracer kinetic measure data in 5-HT targets. N indicates the number of individual participants included in the analysis. R^2^ values are adjusted R^2^.

Using linear models, the observed percent differences per decade were 1.5% in 5-HT-1A receptors, 7.0% in 5-HT-2A receptors, and 3.0% in 5-HT transporters. The rank order of these effects is consistent with the average age correlations yielded from the meta-analyses. These measures are provided as a more concrete quantification of age-related differences relative to 5-HT functioning in younger adulthood.

### 3.5 Statistical power analyses

The average sample sizes of the included studies were about 35 participants (*median* = 22), ranging from 7 to 132 participants. Assuming that the meta-analytic effect sizes closely resembled the true effects of age, we found an average power of 71.0% percent across studies. Half of the studies reached the commonly recommended statistical power of at least 80%. The power analyses revealed the minimum sample sizes required for a minimum of 80% of statistical power in detecting linear effects of age at 146 participants for 5-HT-1A receptors, 13 participants for 5-HT-2A receptors, and 30 participants for 5-HT transporters.

## 4. Discussion

The results of the present meta-analysis of cross-sectional PET and SPECT imaging studies in healthy adults revealed several negative effects of age on serotonergic functioning. We found large negative effects of age in 5-HT-2A receptors (largest in global cortex), moderate negative effects of age in 5-HT transporters (largest in thalamus), and small negative effects of age in 5-HT-1A receptors (largest in parietal cortex). Presynaptic 5-HT-1A autoreceptors in raphe/midbrain, however, were relatively preserved in older age. Overall, the estimated age-related reductions across 5-HT targets ranged between 1.5% and 7.0% per decade. In addition to variation in age effects across 5-HT targets, we identify further variables that moderated age effects. Overall, the study provides the first quantitative synthesis of adult age differences in the 5-HT system.

The meta-analytic evidence for fewer postsynaptic receptors in terminal regions implies a restricted capacity to transmit 5-HT mediated signals in older age. This alteration in signal transduction might be compensated by a mechanism whereby 5-HT could be active in the synaptic cleft for a longer time interval. The activity of synaptic 5-HT could be prolonged due to the lower number of 5-HT transporters with age. The presynaptic transporter proteins may limit clearance of 5-HT from the synaptic cleft via re-uptake into the presynaptic neuron. In contrast to the age effects on post-synaptic receptors in terminal regions, when examining effects within individual brain regions only the negative feedback process that enables the inhibition of 5-HT release into the synaptic cleft seemed to be intact with age. Specifically, there was a non-significant effect of age on 5-HT-1A receptors in raphe/midbrain which are presynaptic somatodendritic autoreceptors. Note that it remains unclear which of these age effects (i.e., pre- versus post-synaptic) appear first and which effects represent only a subsequent auto-regulatory reaction such as triggered by synaptic remodeling (Kitahara et al., 2016; Pineyro and Blier, 1999). Future studies could investigate how age affects other receptor types and presynaptic mechanisms such as synthesis capacity. Unfortunately, there were not enough studies of age differences in synthesis capacity to meta-analyze. One of the few studies on 5-HT synthesis capacity reported stable levels of this presynaptic 5-HT target across the adult life span (Rosa-Neto et al., 2007).

5-HT-1A and 5-HT-2A receptors differ in their distribution across the brain and signaling cascades (Hannon and Hoyer, 2008). For instance, electrophysiological experiments have shown that 5-HT-1A receptor activation induces a hyperpolarization (Nicoll et al., 1990), whereas 5-HT-2A receptor activation primarily leads to a depolarization of the postsynaptic neuron (Aghajanian, 1995). Given the localization of 5-HT-1A and 5-HT-2A on cortical glutamatergic neurons, the relative preservation of 1A compared to 2A receptors might suggest an age-related increase in hyperpolarization of glutamatergic cells. Relatedly, age-related reductions of 5-HT transporters and preservation of somatodendritic 5-HT-1A autoreceptors might predict an age-related increase in 5-HT levels. For a deeper understanding of the functional implications of the present work, the interactions of 5-HT with other neurochemical systems such as glutamate and GABA should be taken into account (Ciranna, 2006).

In general, 5-HT-1A and 5-HT-2A receptors likely mediate distinct neural and behavioral functions (Barnes and Sharp, 1999). A recent theory suggests that 5-HT-1A and 5-HT-2A receptors enable humans to adapt flexibly to adverse events in the environment (Carhart-Harris and Nutt, 2017). Concretely, the mostly inhibitory 5-HT-1A receptor was assumed to mediate passive coping responses such as internal reappraisal of a stressful situation (Carhart-Harris and Nutt, 2017). The mainly excitatory 5-HT-2A receptor however, was suggested to mediate active coping strategies such as engaging in an interpersonal confrontation (Carhart-Harris and Nutt, 2017). In light of this hypothesis, the finding of stronger age-related differences in 5-HT-2A receptors compared to 5-HT-1A receptors might be viewed as consistent with the evidence that older adults use more passive, emotion-focused than active, problem-focused coping strategies (Folkman et al., 1987). The potential mediation of these behavioral age differences by serotonergic function has not been directly tested. Surprisingly, few studies (n=6) examined associations between 5HT receptors and transporters and physiological, emotional, and cognitive variables, primarily focusing on personality traits (Frokjaer et al., 2009; Parsey et al., 2002; Rosell et al., 2010; Zientek et al., 2016) and hormone levels (Frokjaer et al., 2009; Moses-Kolko et al., 2011). Thus, additional research is needed to examine the functional consequences of these neurobiological age differences.

Besides variation across 5-HT targets, we observed that the age effects were moderated by publication year and radionuclide. First, the evidence that smaller effects of age were reported in more recently published studies might be explained by higher effective camera resolutions due to improved technologies and more advanced quantification methods. These newer cameras may better assess individual subregions of the brain and minimize signal mixing with nearby white matter or cerebrospinal fluid (see below for additional discussion of partial volume effects). Unfortunately, we were not able to further investigate this hypothesis since effective scanner resolution is only rarely reported in PET and SPECT imaging studies. Second, the moderating effect of radionuclide might be accounted for by the more reliable and accurate signal of ^18^F because of its longer half-life and shorter positron range compared to ^11^C (Morris et al., 2013). An important caveat here is not to over-interpret the results of the meta-regression. The moderating effects could also be explained by shared confounds between the studies. For instance, we were not able to consider previously reported moderators of the age effect such as body mass index (Hesse et al., 2014), sex hormones (Moses-Kolko et al., 2011), and genes (David et al., 2005). Furthermore, we did not find any evidence for a moderating effect of sex as some studies have reported previously (e.g., Baeken et al., 1998; Cidis Meltzer et al., 2001). Yet, this finding is in line with studies based on larger samples that did also not find an age by sex interaction (e.g., Frokjaer et al., 2009; van Dyck et al., 2000).

A further limitation of the meta-analysis is the potential overestimation of the effects of age. One reason might be publication bias as quantitatively indicated in 5-HT-1A receptors and informally observed during the literature search. A few studies reporting a non-significant age effect did not state the exact size of the age effect and could hence not be included in the meta-analyses. This is a major limitation of the study. Partial volume effects could have additionally contributed to an overestimation of the effects of age in the 5-HT system. There is considerable evidence for grey matter density reductions with age (Sowell et al., 2003). PET and SPECT signals in atrophy-affected, smaller brain regions are more susceptible to blurring by surrounding tissue (Hoffman et al., 1979; Hoffman et al., 1982). Such partial volume effects can lead to an underestimation of the 5-HT signal in age, as has been reported in the dopamine system (Morris et al., 1999; Smith et al., 2017). Several established techniques are available to account for partial volume effects (e.g., Meltzer et al., 1990; Muller-Gartner et al., 1992; Rousset et al., 1998), but only 7 out of 31 included studies applied such corrections. These potential measurement confounds in a majority of the included studies might have led to an overestimation of the age effect in the meta-analyses. Yet, we did not find any evidence for a systematic influence of partial volume corrections on the reported effects of age. To differentiate age-related receptor loss from age-related neuronal atrophy, it will be important for future studies to directly compare age effects in uncorrected data and after correcting for partial volume effects. Additionally, it is important to note that mean levels of binding vary across brain regions, which may bias estimates of age effects. Specifically, age effects may be underestimated due to floor effects in regions that show low levels of binding.

The present study has several additional limitations, both at the level of the meta-analyses and the level of the included studies. First, the literature search was only based on one database as opposed to the recommendations of the PRISMA statement. Yet, we inspected a large number of potentially relevant studies and conducted a thorough reference search. Second, the individual studies seemed to involve on average a lower number of older adults than younger adults as indicated by the scatter plots. This imbalance might have limited the ability to detect nonlinear effects of age on the 5-HT system across adulthood. Third, the individual studies used radionuclides of varying quality. We attempted to include only studies using radionuclides of sufficient affinity and selectivity for 5-HT (Paterson et al., 2013). Finally, our meta-analyses were based on cross-sectional studies which measure age-related *differences* as opposed to age-related *changes* and these age differences may be related to cohort differences rather than developmental change. Both limitations can be addressed with longitudinal study designs. The only longitudinal study so far showed stable 5-HT-2A receptor levels after two years (Marner et al., 2009) but suffered from a small sample size, a limited age range, and a relatively short longitudinal study interval given the estimated rate of decline inferred from the results of cross-sectional studies. Hence, longitudinal studies with sufficient power, a broad age range, and a longer follow-up period will be necessary to disentangle age and cohort effects. Longitudinal studies could further provide insight into the nature of the underlying degradation processes (e.g. zero-order or first-order process), for which the present dataset lacked sufficient granularity.

Here we conducted the first meta-analysis that precisely quantifies adult age differences in several 5-HT targets. The finding of an age-related reduction in 5-HT receptors reveals consistent evidence for limitations in serotonergic signal transmission in older age. Age-related losses in the 5HT system might be compensated by fewer 5-HT transporters, which might lead to a prolonged activity of 5-HT in the synaptic cleft. The regulation of 5-HT release into the synaptic cleft via presynaptic 5-HT-1A autoreceptors may remain intact in older age. The evidence for a relative preservation of 5-HT-1A compared to 5-HT-2A receptors is consistent with a recent theory linking 5-HT functioning to behavioral evidence for age differences in the use of coping strategies. Finally, the present meta-analytic results may also inform clinical studies examining the role of 5-HT in neurological and psychiatric disorders. The results provide an informative baseline for healthy aging to which clinical groups could be compared (e.g., late life depression, Parkinson’s Disease, Alzheimer’s’ Disease). Related to this, the meta-analytic effects were used to estimate required minimum sample sizes in future studies of age differences in 5HT function in healthy adults (5-HT-1A receptors, N=146; 5-HT-2A receptors, N=13; 5-HT transporters, N=30).

## Conflict of interest

The authors declare no competing financial interests.

## Acknowledgements

TMK was supported by the International Research Training Group (IRTG 2150) of the German Research Foundation and by a full doctoral scholarship of the German National Academic foundation. GRSL was supported by US National Institute on Aging Pathway to Independence Award R00-AG042596.

